# Integrated Bioinformatics Analysis of PALB2 Reveals Expression Patterns, Molecular Interactions, and Prognostic Significance in Breast Cancer

**DOI:** 10.64898/2026.07.19.739427

**Authors:** Arifa Jahan Bithi, Mahatab Hasan Rahat

**Affiliations:** Department of Genetic Engineering and Biotechnology, East West University, Dhaka-1212, Bangladesh

**Keywords:** PALB2, Breast Cancer, Bioinformatics, Gene Expression Analysis, Protein–Protein Interaction, Survival Analysis, Mutation Profiling, Homologous Recombination, TCGA, GEPIA2

## Abstract

**Background:** Partner and Localizer of BRCA2 (PALB2) is a key tumor suppressor gene involved in homologous recombination-mediated DNA repair through its interactions with BRCA1 and BRCA2. Germline alterations in PALB2 have been associated with hereditary breast cancer risk; however, its broader molecular role in breast cancer progression and prognosis requires further investigation.

**Methods:** A comprehensive in-silico analysis of PALB2 was performed using publicly available databases and bioinformatics platforms. Differential expression of PALB2 in breast cancer were evaluated using GEPIA2. Prognostic significance was assessed through Kaplan–Meier analyses for overall survival (OS) and disease-free survival (DFS). Protein–protein interaction (PPI) networks were constructed using STRING. Functional enrichment analyses of PALB2-associated genes were conducted using g. Mutational profiling of PALB2 in breast cancer was performed using cBioPortal with data from TCGA breast cancer cohorts.

**Results:** PALB2 expression was elevated in breast tumor tissues compared with normal breast tissues. Survival analyses revealed no statistically significant association between PALB2 expression and either overall survival (HR = 0.88, p = 0.44) or disease-free survival (HR = 0.74, p = 0.11). Protein interaction analysis revealed strong interactions between PALB2 and major DNA repair proteins including BRCA1, BRCA2, RAD51, RAD51C, FANCD2, and BRIP1. Functional enrichment analysis showed limited significant pathway enrichment, with only marginal transcription factor motif enrichment observed. Mutational analysis demonstrated diverse genomic alterations including missense mutations, truncating mutations, copy number gains, and shallow deletions.

**Conclusion:** The findings support the biological relevance of PALB2 in breast cancer through its elevated expression and strong connectivity within DNA repair pathways. However, PALB2 expression alone does not appear to serve as an independent prognostic indicator. Further studies integrating genomic, transcriptomic, and clinical parameters are required to clarify its role in breast cancer progression and therapeutic response.

## 1. Introduction

Breast cancer remains the most frequently diagnosed malignancy among women and continues to be a leading cause of cancer-related mortality worldwide. Despite substantial advances in diagnosis and treatment, breast cancer continues to pose a major public health challenge, particularly in low- and middle-income countries where delayed diagnosis and limited access to molecular screening strategies contribute to poor clinical outcomes. The development and progression of breast cancer involve complex interactions between environmental exposure, lifestyle factors, and inherited genetic susceptibility (Lichtenstein et al., 2000).

Genetic alterations affecting DNA repair pathways play a critical role in breast cancer development. Among susceptibility genes, Partner and Localizer of BRCA2 (PALB2) has emerged as an important tumor suppressor due to its essential role in maintaining genomic stability. PALB2 functions as a molecular bridge connecting BRCA1 and BRCA2 and facilitates homologous recombination-mediated repair of DNA double-strand breaks. Defects in this repair mechanism can lead to genomic instability and accumulation of mutations that contribute to tumor initiation and progression (Xia et al., 2006; Wu et al., 2020).

The clinical importance of PALB2 has become increasingly recognized over recent years. Germline pathogenic variants in PALB2 have been associated with substantially elevated breast cancer risk and are now routinely included in multigene cancer susceptibility testing panels. Recent evidence further supports PALB2 as a moderate-to-high penetrance breast cancer susceptibility gene, with mutation carriers exhibiting significantly increased lifetime breast cancer risk (Ruberu et al., 2024).

Beyond inherited pathogenic variants, increasing attention has been directed toward understanding the broader biological functions of PALB2 in breast cancer progression. Previous studies have suggested that altered PALB2 activity may influence tumor development through modulation of DNA repair efficiency, maintenance of chromosomal integrity, and therapeutic response mechanisms (Wu et al., 2020). Furthermore, emerging molecular studies continue to identify PALB2-associated genomic alterations across diverse breast cancer subtypes, highlighting its potential clinical relevance in precision oncology and targeted therapeutic strategies.

The growing availability of large-scale genomic databases and computational platforms has enabled comprehensive investigations into the molecular behavior of cancer-associated genes. Integrated bioinformatics approaches allow characterization of gene expression patterns, prognostic significance, protein interaction networks, pathway enrichment, and mutational landscapes using publicly available datasets. Such approaches can provide important biological insights and generate hypotheses for future experimental validation.

Therefore, the present study aimed to perform a comprehensive in-silico characterization of PALB2 in breast cancer through integrated bioinformatics analyses, including expression profiling, survival assessment, protein–protein interaction analysis, functional enrichment evaluation, and mutational profiling to better understand its molecular significance in breast cancer biology.

## 2. Materials and Methods

### 2.1 Tissue Expression Profile Analysis of PALB2

To evaluate the baseline expression pattern of *PALB2* across normal human tissues, tissue-specific gene expression data were obtained from the <u>GTEx Portal</u>. Expression levels of *PALB2* across multiple human tissue types were assessed using transcript abundance values represented as Transcripts Per Million (TPM). Distribution patterns across tissues were visualized using violin plots to compare relative expression levels and variability among tissue types.

### 2.2 Differential Expression Analysis

Differential expression analysis of PALB2 in breast cancer was performed using the Gene Expression Profiling Interactive Analysis version 2 (GEPIA2) web server. The “Expression DIY” module was used to compare PALB2 expression levels between breast invasive carcinoma (BRCA) tissues and normal breast tissues. Expression values were represented as log2(TPM+1).

### 2.3 Survival Analysis

The prognostic significance of PALB2 expression was evaluated using GEPIA2. Kaplan–Meier survival analyses were performed for both overall survival (OS) and disease-free survival (DFS) among breast cancer patients. Hazard ratios and log-rank p-values were used for statistical interpretation.

### 2.4 Protein–Protein Interaction Network Analysis

Protein interaction analysis was performed using the STRING database. PALB2 and associated genes involved in DNA repair mechanisms were analyzed, and interactions with confidence scores greater than 0.7 were retained for network construction.

### 2.5 Functional Enrichment Analysis

Functional enrichment analysis of PALB2 and interacting genes was conducted using g. Gene Ontology (GO) categories, Kyoto Encyclopedia of Genes and Genomes (KEGG), Reactome pathways, and transcription factor motifs were evaluated.

### 2.6 Mutation Analysis

Mutational profiling of PALB2 was conducted using cBioPortal with breast cancer datasets from The Cancer Genome Atlas (TCGA). Mutation types and frequencies were evaluated to characterize the genomic landscape of PALB2 in breast cancer.

## 3. Results

### 3.1 Tissue-Specific Expression Pattern of PALB2

The tissue expression profile analysis demonstrated that *PALB2* exhibits variable expression across different human tissues. Higher expression levels were observed in tissues with increased proliferative activity and metabolic demand, including cultured fibroblasts, transformed lymphocytes, and several epithelial tissues. Moderate expression was detected in tissues such as breast, lung, and reproductive tissues, whereas relatively lower expression levels were observed in tissues including whole blood and certain brain regions.

**Figure 01:**
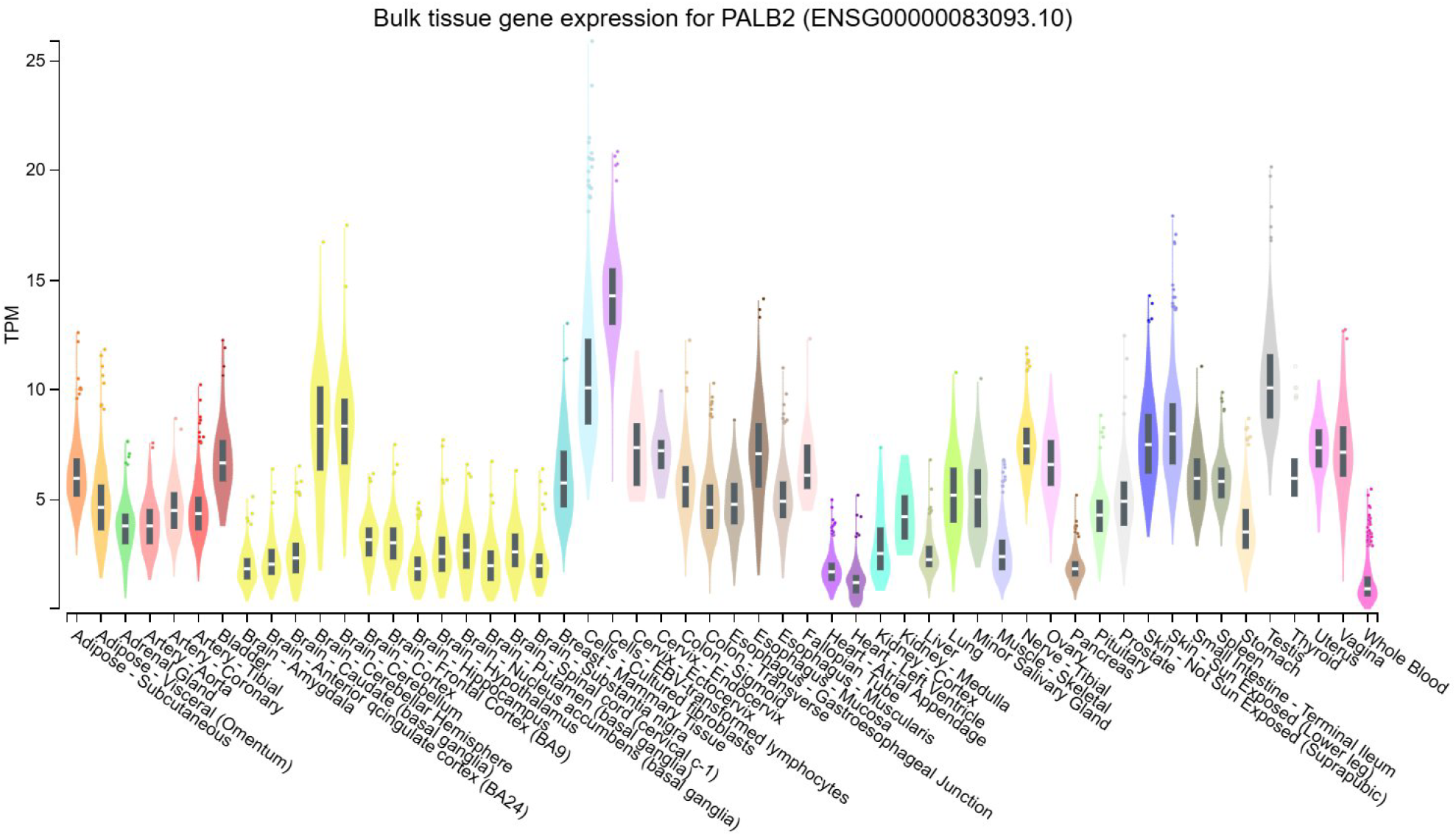
Normal Expression of PALB2 Across the Tissue

The broad distribution of *PALB2* expression across multiple tissues is consistent with its fundamental biological role in maintaining genomic integrity through DNA repair mechanisms. The observed expression pattern suggests that *PALB2* functions as a widely expressed housekeeping-associated gene involved in cellular maintenance processes rather than displaying highly tissue-specific expression. Additionally, detectable expression in breast tissue supports its biological relevance in breast cancer development and justifies further investigation of its molecular characteristics in breast tumors.

### 3.2 PALB2 Expression in Breast Cancer

Analysis of PALB2 expression demonstrated increased expression levels in breast tumor tissues compared with normal breast tissues. Tumor samples showed a generally higher distribution of expression values, indicating upregulation of PALB2 in breast cancer tissues.

**Figure 02:**
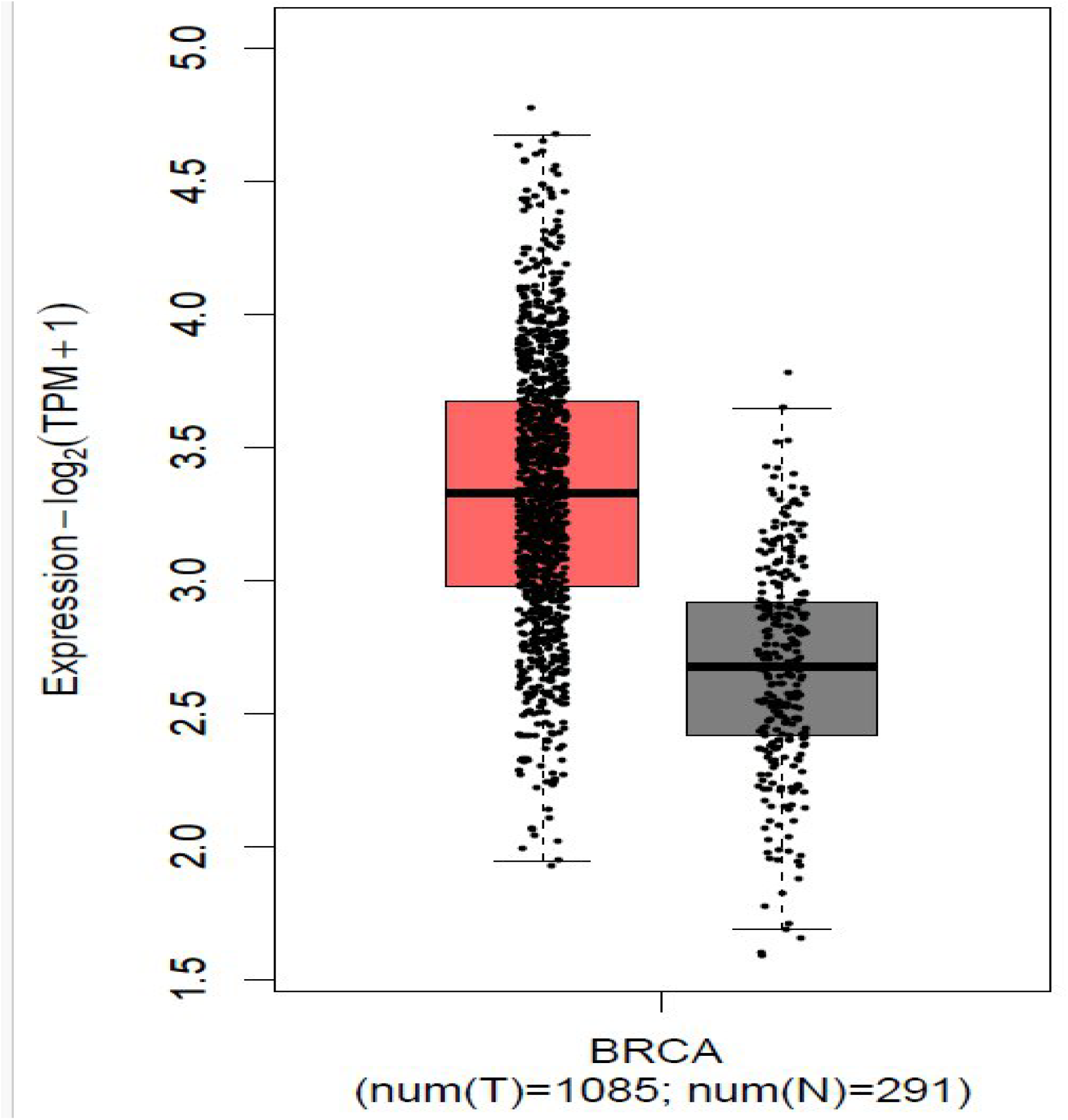
PALB2 Expression Levels in Breast Tumor and Normal Tissues.

### 3.3 Prognostic Significance of PALB2 Expression

Kaplan–Meier overall survival analysis demonstrated no statistically significant association between PALB2 expression and patient survival (HR=0.88, p=0.44). Survival curves between high- and low-expression groups remained closely aligned.

**Figure 03:**
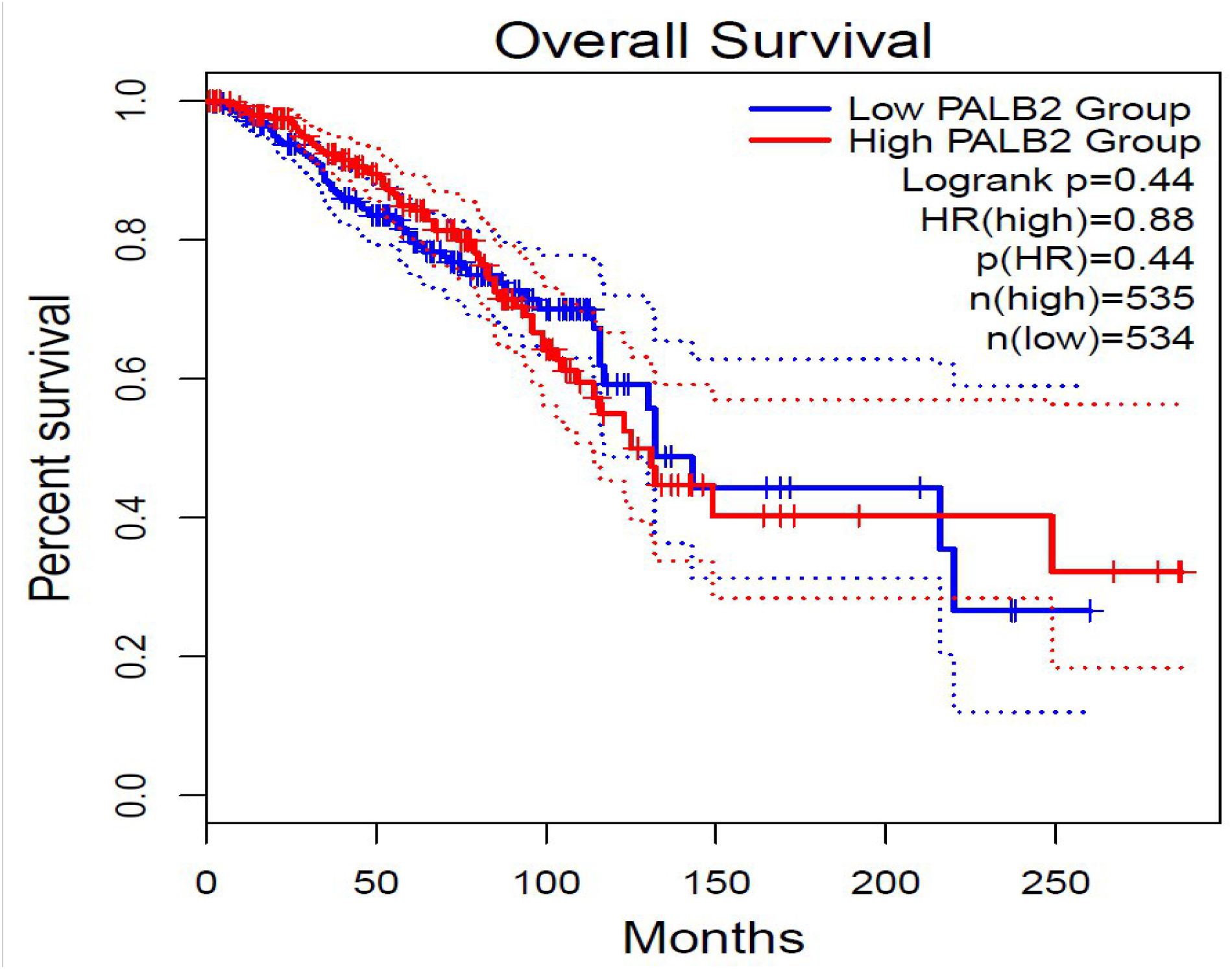
Kaplan–Meier Overall Survival Curve Based on PALB2 Expression in Breast Cancer Patients.

Similarly, disease-free survival analysis revealed no statistically significant differences between expression groups (HR=0.74, p=0.11). Although patients with elevated PALB2 expression showed a slightly improved trend in recurrence-free survival, the observed difference was not statistically significant.

**Figure 04:**
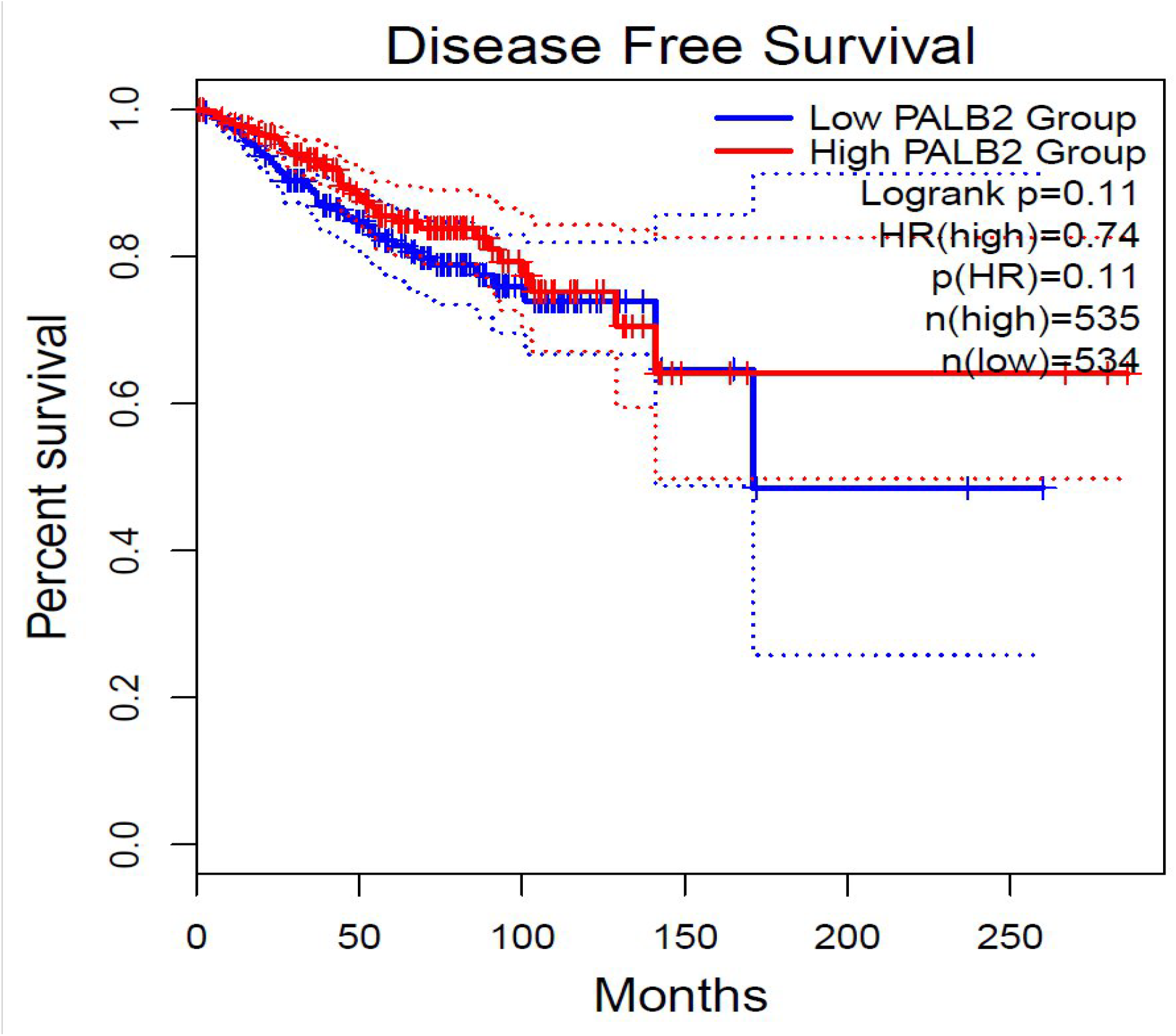
Kaplan–Meier Disease-Free Survival Curve Based on PALB2 Expression in Breast Cancer Patients.

### 3.4 Protein–Protein Interaction Network

The STRING analysis identified multiple high-confidence interactions between PALB2 and genes involved in DNA repair pathways. Major interacting proteins included BRCA1, BRCA2, RAD51, RAD51C, FANCD2, BRIP1, and MORF4L1. The network demonstrated extensive connectivity among proteins involved in homologous recombination repair mechanisms.

**Figure 05:**
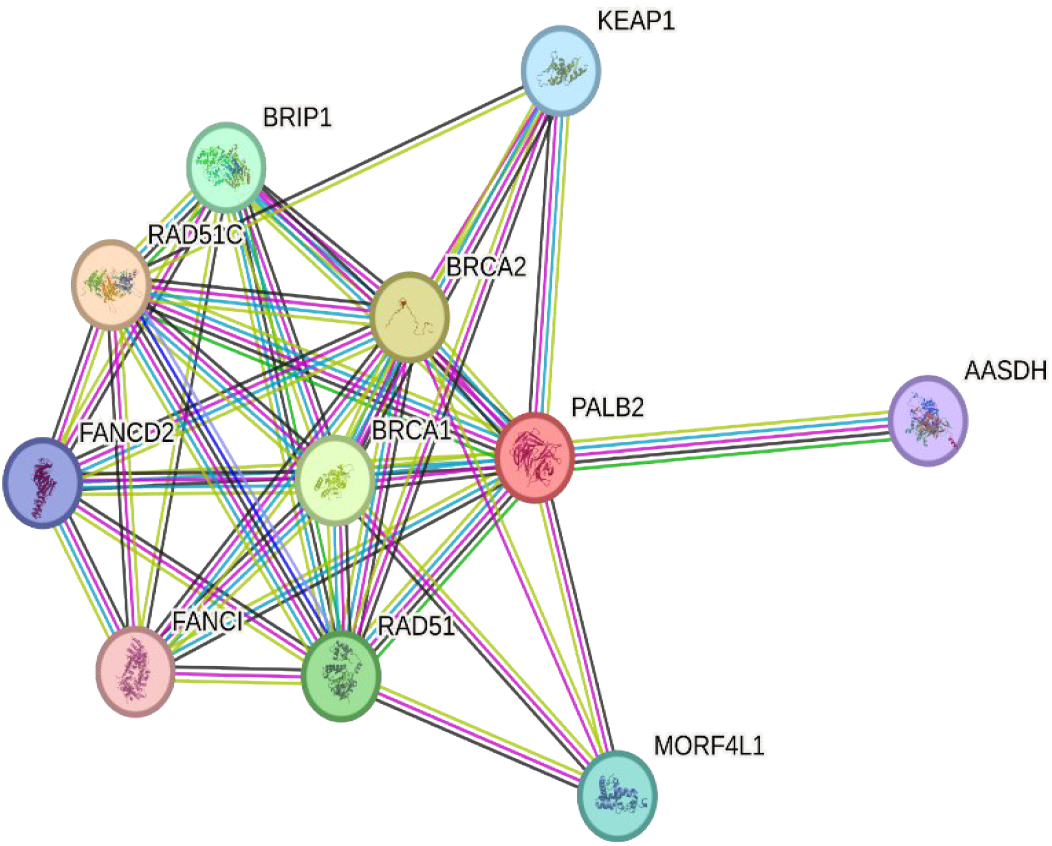
Protein–protein interaction network of PALB2 and associated genes generated using STRING.

### 3.5 Functional Enrichment Analysis

Functional enrichment analysis showed no statistically significant enrichment for major biological processes or signaling pathways following multiple testing correction. A transcription factor motif associated with ZNF772 demonstrated marginal significance, whereas broader pathway-level associations were not observed.

**Figure 06:**
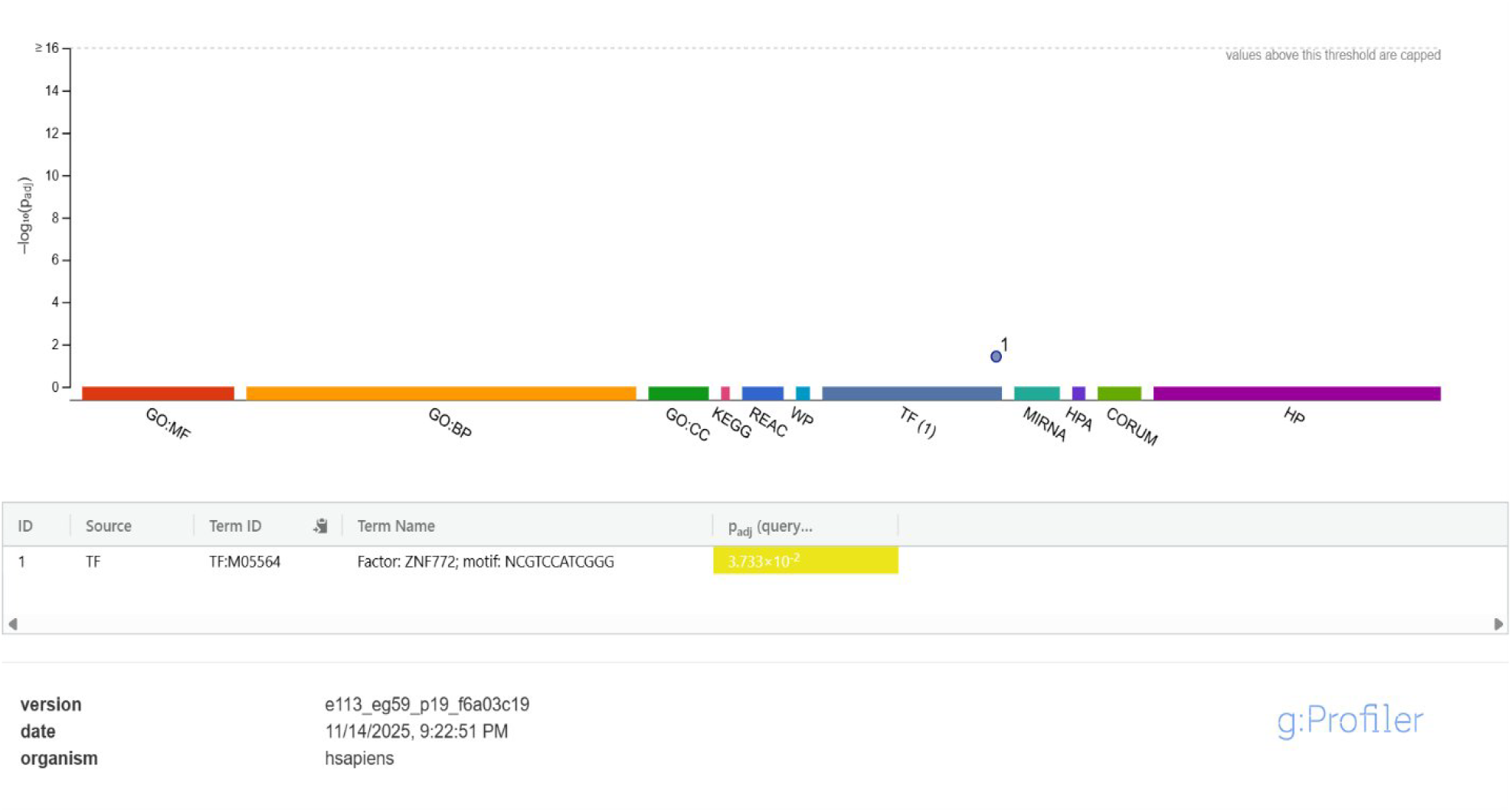
Functional enrichment analysis of PALB2-associated genes generated using g:Profiler.

### 3.6 Mutational Profile of PALB2

Mutation analysis demonstrated several genomic alterations in PALB2, including missense mutations, truncating mutations, copy number gains, amplifications, and shallow deletions. Mutation events were distributed across multiple samples without evidence of a dominant hotspot region.

**Figure 07:**
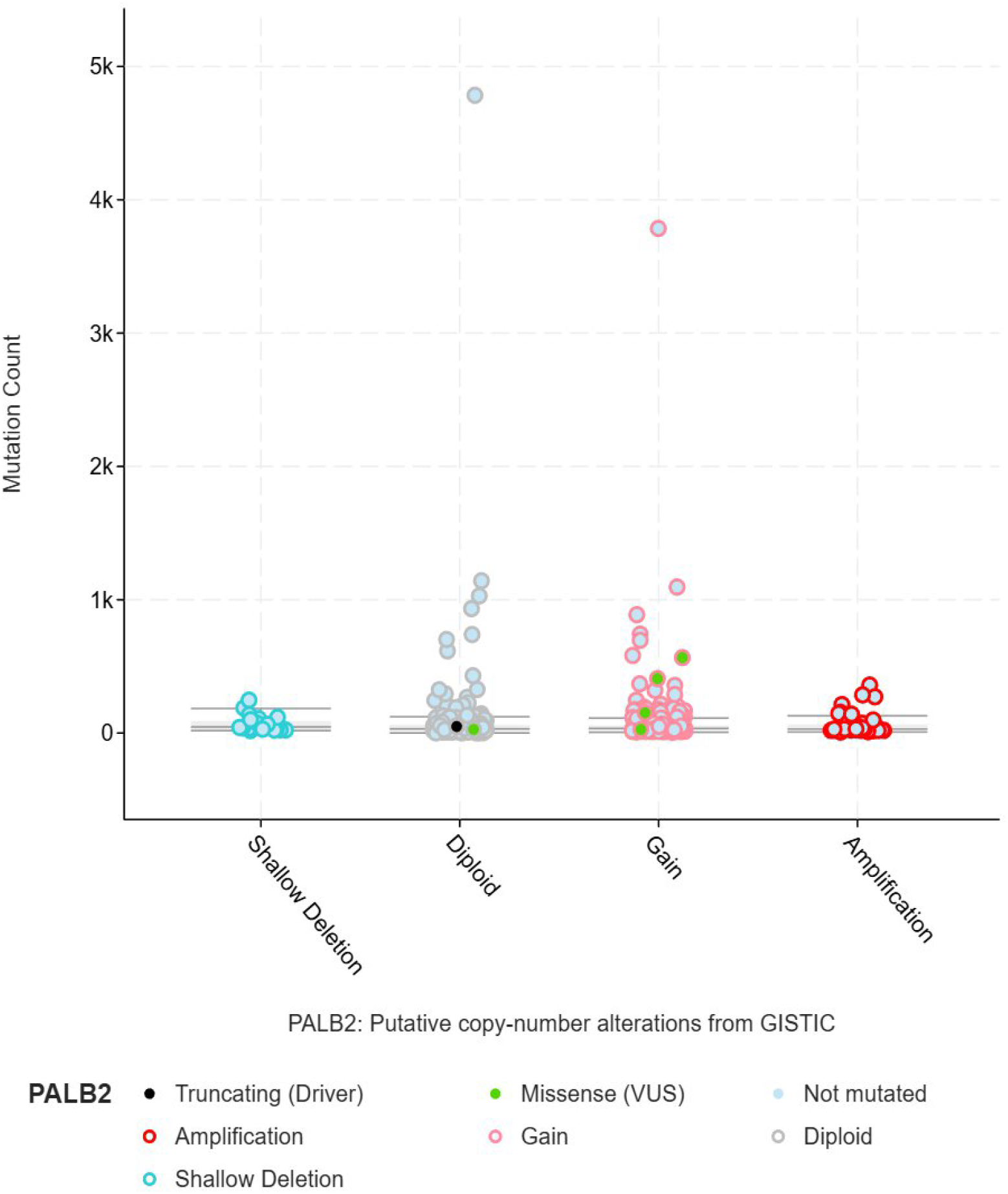
Distribution of PALB2 mutations and copy-number alterations in the TCGA breast cancer cohort.

## 4. Discussion

The present study performed an integrated bioinformatics evaluation of *PALB2* to investigate its molecular characteristics and potential significance in breast cancer. Tissue-specific expression profiling demonstrated that *PALB2* is broadly expressed across multiple normal human tissues, although expression levels varied considerably among tissue types. Elevated expression in proliferative tissues and transformed cell populations may reflect the fundamental requirement for efficient DNA repair mechanisms in actively dividing cells. The widespread expression pattern supports the concept that *PALB2* functions as a genome maintenance-associated gene involved in essential cellular processes rather than as a highly tissue-specific protein. Furthermore, detectable expression in normal breast tissue reinforces its biological relevance in breast tissue homeostasis and carcinogenesis.

The differential expression analysis further revealed increased *PALB2* expression in breast tumor tissues compared with normal breast tissues, suggesting a potential involvement in tumor-associated cellular processes and DNA damage response mechanisms. The increased expression observed in tumors may represent an adaptive cellular response to replication stress and genomic instability, which are characteristic features of malignant cells. Cancer cells frequently accumulate DNA damage during uncontrolled proliferation, and increased expression of homologous recombination-associated genes may serve as a compensatory mechanism for maintaining genomic integrity and promoting tumor cell survival.

Despite the observed differential expression between tumor and normal tissues, survival analyses demonstrated no statistically significant association between *PALB2* expression and either overall survival or disease-free survival. Although a modest trend toward improved disease-free survival was observed in patients with higher *PALB2* expression, the lack of statistical significance suggests that expression level alone may not function as an independent prognostic biomarker. Breast cancer outcomes are strongly influenced by multiple clinical and molecular variables, including tumor subtype, histological grade, treatment strategy, and genomic background, which may mask the independent contribution of *PALB2* expression.

Protein interaction analysis reinforced the central role of *PALB2* within homologous recombination-mediated DNA repair pathways. Strong interactions observed with *BRCA1, BRCA2, RAD51, FANCD2*, and related proteins support previously established evidence describing *PALB2* as a molecular scaffold coordinating DNA repair machinery. These interactions highlight the importance of *PALB2* in maintaining genome stability and regulating cellular responses to DNA damage.

Interestingly, functional enrichment analysis did not identify strong pathway-level significance despite the known biological importance of *PALB2*. This finding may reflect the structural and interaction-dependent role of *PALB2*, where biological function is mediated primarily through molecular partnerships rather than independent pathway activity. In addition, statistical correction methods applied during enrichment analysis may reduce significance for genes already represented within highly curated DNA repair pathways.

Mutation profiling further demonstrated heterogeneous genomic alterations within *PALB2*, including missense mutations, truncating variants, copy number gains, and deletions. Truncating mutations may disrupt essential interaction domains required for homologous recombination repair and potentially contribute to homologous recombination deficiency, whereas copy number alterations may influence expression levels and functional activity.

Overall, the findings suggest that PALB2 plays a role in molecular pathways linked to breast cancer development and DNA repair processes, but its prognostic utility may be limited when evaluated independently. Future investigations integrating molecular subtype-specific analyses, larger datasets, and functional experimental validation are needed to clarify the precise clinical significance of *PALB2* in breast cancer.

## 5. Conclusion

The present study conducted an integrated bioinformatics analysis to investigate the molecular characteristics and potential clinical significance of *PALB2* in breast cancer. Tissue-specific expression analysis demonstrated that *PALB2* is broadly expressed across multiple human tissues, supporting its role as an essential gene involved in genome maintenance and DNA repair processes. Detectable expression in breast tissue further supports its biological relevance in breast cancer research.

Differential expression analysis demonstrated increased *PALB2* expression in breast tumor tissues compared with normal breast tissues, suggesting involvement in cancer-associated biological processes and DNA damage response mechanisms. However, despite elevated expression levels, survival analyses revealed no statistically significant association with overall or disease-free survival, indicating that *PALB2* expression alone may not serve as a reliable prognostic marker.

Protein–protein interaction analysis identified strong associations between *PALB2* and major DNA repair proteins, including *BRCA1, BRCA2*, and *RAD51*, confirming its central role in homologous recombination pathways and genome stability maintenance. Furthermore, mutation profiling revealed diverse genomic alterations, including missense mutations, truncating mutations, and copy number variations, suggesting heterogeneous mechanisms through which *PALB2* may contribute to breast cancer biology.

Collectively, these findings highlight the biological importance of *PALB2* in breast cancer development and DNA repair regulation while suggesting limited independent prognostic value.

Further studies involving larger datasets, subtype-specific analyses, and experimental validation are required to better understand the functional and clinical implications of *PALB2* in breast cancer.

## Disclaimer/Publisher’s Note

The statements, opinions and data contained in all publications are solely those of the individual author(s) and contributor(s) and not of MDPI and/or the editor(s). MDPI and/or the editor(s) disclaim responsibility for any injury to people or property resulting from any ideas, methods, instructions or products referred to in the content.

